# Intrinsic resistance networks shape cefiderocol susceptibility in ST258 *Klebsiella pneumoniae*

**DOI:** 10.64898/2026.02.12.705544

**Authors:** Kevin J Rome, Austin J. Terlecky, Kelly K. Yen, Tengfei Long, Mia Bucich, Klara V. Thom, Elena Shashkina, Liang Chen, Barry Kreiswirth

## Abstract

Cefiderocol (CFDC) is a siderophore-conjugated cephalosporin developed to overcome multidrug resistance in Gram-negative bacteria. Despite its unique iron-dependent entry mechanism, CFDC resistance has emerged in *Klebsiella pneumoniae*, primarily driven by alterations in siderophore transport and β-lactamase evolution; however, the broader intrinsic resistome that supports CFDC tolerance remains incompletely defined. Here, we performed high-density transposon mutagenesis (Tn-Seq) in the epidemic ST258 *K. pneumoniae* to map the functional genetic landscape of CFDC susceptibility. Tn-Seq identified siderophore uptake components (*tonB* and *cirA*) as the dominant determinants of CFDC resistance. In contrast, disruption of genes involved in peptidoglycan recycling (*ampG, ldcA*), synthesis (*mrcB, lpoB*) and enterobacterial common antigen (ECA) biosynthesis (*wzxE, wzyE*), as well as deletion of plasmid-encoded *bla*_KPC-3_, increased CFDC susceptibility. In a CFDC-resistant Δ*tonB* strain, targeting these envelope homeostasis pathways yielded only limited resensitization relative to the siderophore-competent parental strain. Deletion of *bla*_KPC-3_ produced the greatest increase in susceptibility, reducing the CFDC MIC four-fold. This pattern is consistent with a model in which reduced CFDC influx in the Δ*tonB* background lowers intracellular drug exposure to levels at which the otherwise limited anti-CFDC activity of KPC β-lactamase becomes sufficient to drive resistance. Together, these data define a hierarchical genetic architecture for CFDC resistance in ST258 *K. pneumoniae*, in which iron-dependent drug uptake is primary, β-lactamase activity is secondary, and intrinsic envelope stress buffering shapes bacterial fitness once CFDC enters the cell.

## Introduction

Carbapenem-resistant *Klebsiella pneumoniae* (CRKP) remains a preeminent global public health threat, characterized by high mortality rates and a diminishing arsenal of effective therapeutics (1–4). Within this group, the sequence type 258 (ST258) lineage is of particular clinical concern; it is the dominant carbapenemase-producing clone worldwide, primarily driven by the dissemination of *K. pneumoniae* carbapenemases (KPCs) (5–8). While recent years have seen the introduction of newer β-lactam/β-lactamase inhibitor (BL/BLI) combinations such as ceftazidime-avibactam, their utility is increasingly restricted by the rapid emergence of resistance, most notably through *bla*_KPC_ mutations and significant clinical toxicities associated with older “last-resort” agents like colistin (9–11).

To address this therapeutic gap, cefiderocol (CFDC), a first-in-class siderophore-conjugated cephalosporin, was recently developed (12–15). CFDC utilizes a unique “Trojan horse” mechanism to bypass the permeability barriers of Gram-negative bacteria (16). The molecule consists of a cephalosporin core linked to a catechol moiety that chelates extracellular ferric iron (Fe^3+^)(17). This complex is then actively imported into the periplasm via TonB-dependent transporters (TBDTs)-such as CirA and Fiu in *K. pneumoniae*-thereby exploiting the bacterial necessity for iron acquisition (18–21). Once in the periplasm, CFDC binds with high affinity to penicillin-binding proteins (PBPs), primarily PBP3, to inhibit peptidoglycan synthesis (22,23).

Despite its potent activity against multi-drug resistant (MDR) isolates, clinical resistance to CFDC has emerged rapidly in several pathogenic species, including *K. pneumoniae, Acinetobacter baumannii*, and *Pseudomonas aeruginosa* (21,24–27). Current knowledge suggests that CFDC resistance is predominantly mediated by two distinct axes: (i) impairment of the siderophore-mediated uptake machinery, often involving mutations in the *tonB* complex or specific TBDTs like *cirA* (18–20,28,29), and (ii) high-level expression or structural evolution of β-lactamases. Specifically, mutations in *bla*_KPC-3_ leading to variants such as KPC-31, as well as the presence of NDM-type metallo-β-lactamases, have been shown to significantly elevate CFDC MICs (13,30–33).

While these primary mechanisms are well-documented, emerging evidence indicates that CFDC susceptibility is a complex, polygenic trait (25,34–36). Recent studies have suggested that the “resistome” extends beyond simple uptake and hydrolysis, involving broader physiological pathways such as cell envelope stress responses and oxidative stress mitigation (25,34,35). However, a comprehensive, genome-wide investigation into the fitness determinants that govern both resistance and the potential for resensitization in a clinically relevant ST258 background is currently lacking.

In this study, we utilized high-density transposon mutagenesis (Tn-Seq) to define the functional genomic landscape of CFDC susceptibility in the ST258 clinical isolate BK30684 (37). This methodology enabled us to confirm established resistance mechanisms while identifying a suite of underappreciated determinants, including those involved in nitrogen metabolism, oxidative stress, and plasmid-encoded loci. Furthermore, using CRISPR-Cas9-based gene editing and plasmid curing, we dissected the hierarchical architecture of these resistance layers. Our findings reveal that while TonB-mediated uptake remains the dominant factor, specific envelope-associated pathways constitute a secondary layer of resistance that can be exploited to resensitize MDR strains. This work provides a comprehensive functional genomics map of CFDC fitness determinants, offering a roadmap for preserving the efficacy of this vital antibiotic against MDR pathogens.

## Material and Methods

### Bacterial strains and growth conditions

All bacterial strains and plasmids used in this study are listed in Table S1. *K*.*pneumoniae* strains were routinely cultured in Luria-Bertani (LB) broth or on LB agar plates at 37°C. For CFDC susceptibility testing, strains were grown on cation-adjusted Mueller-Hinton agar (CAMHA) or in iron-depleted cation-adjusted Mueller-Hinton broth (IDMH), as specified. IDMH was prepared by treating CAMHB with Chelex-100 resin (Bio-Rad) for 6 hours to remove divalent cations, followed by re-supplementation with Ca^2+^, Mg^2+^, and Zn^2+^ to the concentrations recommended by the Clinical and Laboratory Standards Institute (CLSI) (33). All *K*.*pneumoniae* strains were derived from clinical isolate BK30684 (39). *Escherichia coli* K-12 strain DH10B was used for conjugative plasmid transfer into *K*.*pneumoniae*. Antibiotic selection was tested as follows: apramycin at 30 μg/mL; rifampin at 100 μg/mL and chloramphenicol at 50μg/mL.

### Antibiotic susceptibility testing

Minimum inhibitory concentrations (MICs) were determined using the broth microdilution method according to CLSI guidelines. For each assay, overnight bacterial cultures were diluted to approximately 5 × 10^5^ CFU/mL in the appropriate medium and incubated with serial two-fold dilutions of the antibiotic in 96-well microtiter plates at 37°C for 16–20 hours. The MIC was defined as the lowest concentration that prevented visible bacterial growth. Reported MIC values were reproducible across a minimum of three independent biological replicates.

### Transposon mutagenesis

A transposon (Tn) mutant library was generated in the ST258 *K. pneumoniae* strain BK30684 using the *Himar*1 mariner transposon carried on the conjugative plasmid pSAMtac1, as previously described (40). Briefly, pSAMtac1, which encodes Himar1 with an apramycin (APR) resistance cassette, was mobilized into BK30684 from the diaminopimelic acid (DAP)-auxotrophic *E. coli* donor WM3064 by conjugation. Following overnight mating, transconjugants were selected on LB agar supplemented with apramycin (30 μg/mL) and IPTG (0.1 mM), and incubated at 37°C. Apramycin-resistant colonies were pooled, resuspended in LB broth with 30% glycerol, and stored at –80°C as an ST258 transposon library.

### Growth Kinetic Assay

Growth kinetic of *K. pneumoniae* BK30684 and its knockout derivatives were assessed in iron-depleted Mueller–Hinton broth (IDMH). Overnight cultures were diluted to approximately 5 × 10^5^ CFU/mL in 200 μL of IDMH, supplemented with or without CFDC at 0.25× the MIC of BK30684, and dispensed into 96-well microtiter plates (Costar). Plates were incubated at 37 °C with continuous shaking, and OD_600_ measurements were recorded every 20 min for 16 h using a multifunctional microplate reader (Tecan Infinite 200 Pro). Each condition was tested in three biological replicates. Growth curves were generated from mean OD_600_ values, and variability between biological replicates was reported as standard deviation.

### CFDC Challenge of Transposon Mutant Libraries

An aliquot of the transposon mutant library was thawed and grown overnight at 37°C in 50 mL LB broth supplemented with apramycin (30 μg/mL). Cultures were diluted to an initial OD_600_ of 0.05 in iron-depleted Mueller–Hinton (IDMH) broth with or without CFDC (4 μg/mL), and incubated at 37°C until reaching an OD_600_ of ~1.0. Cells were harvested for genomic DNA extraction and transposon junction sequencing. Three biological replicates were performed for each condition.

### Deep sequencing of Tn insertions and analysis of sequencing data

Genomic DNA was extracted using the Promega SV Wizard Genomic DNA Purification Kit, and Tn-adjacent fragments were prepared with the NEBNext Ultra II FS DNA Library Prep Kit (New England Biolabs), with minor modifications to published protocols (40). Libraries were sequenced on an Illumina HiSeq platform, generating 10.4 million 150-bp paired-end reads. Reads were trimmed for quality and adapter sequences with Trimmomatic v0.39 (41) and processed using the Tn-Seq PreProcessor (TPP) (42) to quantify insertions at TA dinucleotide sites in the *K. pneumoniae* ST258 strain BK30684 reference genome (GenBank accession NZ_CP006918.1). Gene essentiality was inferred using Hidden Markov Model (HMM) implemented in TRANSIT, which estimates the probability of essentiality based on read counts and the surrounding distribution of TA sites (42,43). Genes with log_2_-fold-change |log_2_FC| > 1 and *q* < 0.05 were identified as CFDC-susceptibility or resistance determinants.

### Gene knockout

Knockout mutants of *K. pneumoniae* BK30684 were constructed using a CRISPR-Cas9 system as described previously (44). Briefly, the parental strain was first transformed with the Cas9/λ-Red helper plasmid pCasKP and selected on LB agar containing apramycin (30 μg/mL). For editing, arabinose-induced recipient cells harboring the Cas9 plasmid were co-electroporated with a spacer-introduced sgRNA plasmid (pSGKP) and a donor repair template, which consisted of either (i) a double-stranded DNA fragment with flanking homology arms and a chloramphenicol resistance cassette or (ii) a 90-nt single-stranded oligonucleotide. Transformants were plated on LB agar supplemented with apramycin (30 μg/mL), rifampin (100 μg/mL), and, when required, chloramphenicol (50 μg/mL), and incubated at 37 °C overnight. Successful edits were confirmed by PCR and Sanger sequencing, and editing plasmids were cured by *sacB*-mediated counterselection.

### Plasmid curing

To eliminate pNJST258C1 from ST258 CRKP strain BK30684, we employed the CRISPR-Cas based vector pLCasCureT as previously described. This construct harbors a guide RNA targeting selected plasmid replicons and encodes Cas9 under the control of the arabinose-inducible pBAD promoter. The vector was introduced into the parental isolate by conjugation, after which Cas9 expression was induced. Transconjugants were screened for plasmid loss by PCR targeting the replicon sequence. Following confirmation of plasmid curing, pLCasCureT was removed through *sacB*-mediated counterselection. The final mutant strain retained an unaltered chromosome and was cured of plasmids pNJST258C1 (45).

## Results

### Transposon mutagenesis of *Klebsiella pneumoniae* ST258 CRKP

We constructed a Himar1 transposon mutant library in *Klebsiella pneumoniae* ST258 carbapenem-resistant strain BK30684. This strain contains a 5.29 Mbp chromosome and three plasmids: pNJST258C1 (86 kb), pNJST258C2 (25.8 kb), and pNJST258C3 (12.4 kb) (6). BK30684 harbors chromosomal *bla*_SHV-11_ and plasmid-borne *bla*_KPC-3_ (pNJST258C2) and *bla*_TEM_ (pNJST258C1), conferring resistance to most β-lactams (Table S3), including carbapenems (imipenem, meropenem), cephalosporins (ceftazidime, cefepime, ceftriaxone), and aztreonam. However, the strain remains susceptible to ceftazidime-avibactam (CAZ/AVI) and CFDC.

To investigate genetic determinants of CFDC susceptibility, we generated a saturated Himar 1 transposon mutant library and performed Illumina-based Tn-Seq after growth in IDMH broth. Median insertion saturation was 52% across chromosomal TA sites, and 62%, 74%, and 62% for plasmids pNJST258C1, pNJST258C2, and pNJST258C3, respectively. These values exceed the ~30% saturation threshold required for robust statistical analyses, including Hidden Markov Model (HMM) classification of essential genes, resampling, and ANOVA (43). Using HMM analysis, we classified chromosomal genes into 491 essential, 4,379 non-essential, 210 growth-defect, and 45 growth-advantage loci (Figure 1A; Table S3). The number of essential genes identified is consistent with previous findings for ST258 *K. pneumoniae* (46). Plasmid-level HMM analysis revealed locus-dependent underrepresentation of Himar1 insertions (Figure 1B; Table S3). In pNJST258C1 and pNJST258C2, replication-associated genes were strongly underrepresented, with no insertions detected in *repA* (pNJST258C1) or RS31270 (encoding a replication initiation protein, pNJST258C2). Reduced insertion densities were also observed in genes involved in plasmid replication and conjugation (*tra* operon, *finO*, and the *rep* operon in pNJST258C1). Disruption of these genes likely results in plasmid loss. Additional underrepresented loci included RS26765 (cloacin immunity family protein, pNJST258C2), RS27065 (antirestriction protein, pNJST258C1), and several hypothetical proteins– encoding genes on pNJST258C1 (RS27230, RS27155, RS27125 and RS27130). Together, these data indicate that replication and partitioning functions are critical for plasmid maintenance, while other underrepresented loci may contribute to plasmid stability or host fitness.

**Figure 1.**
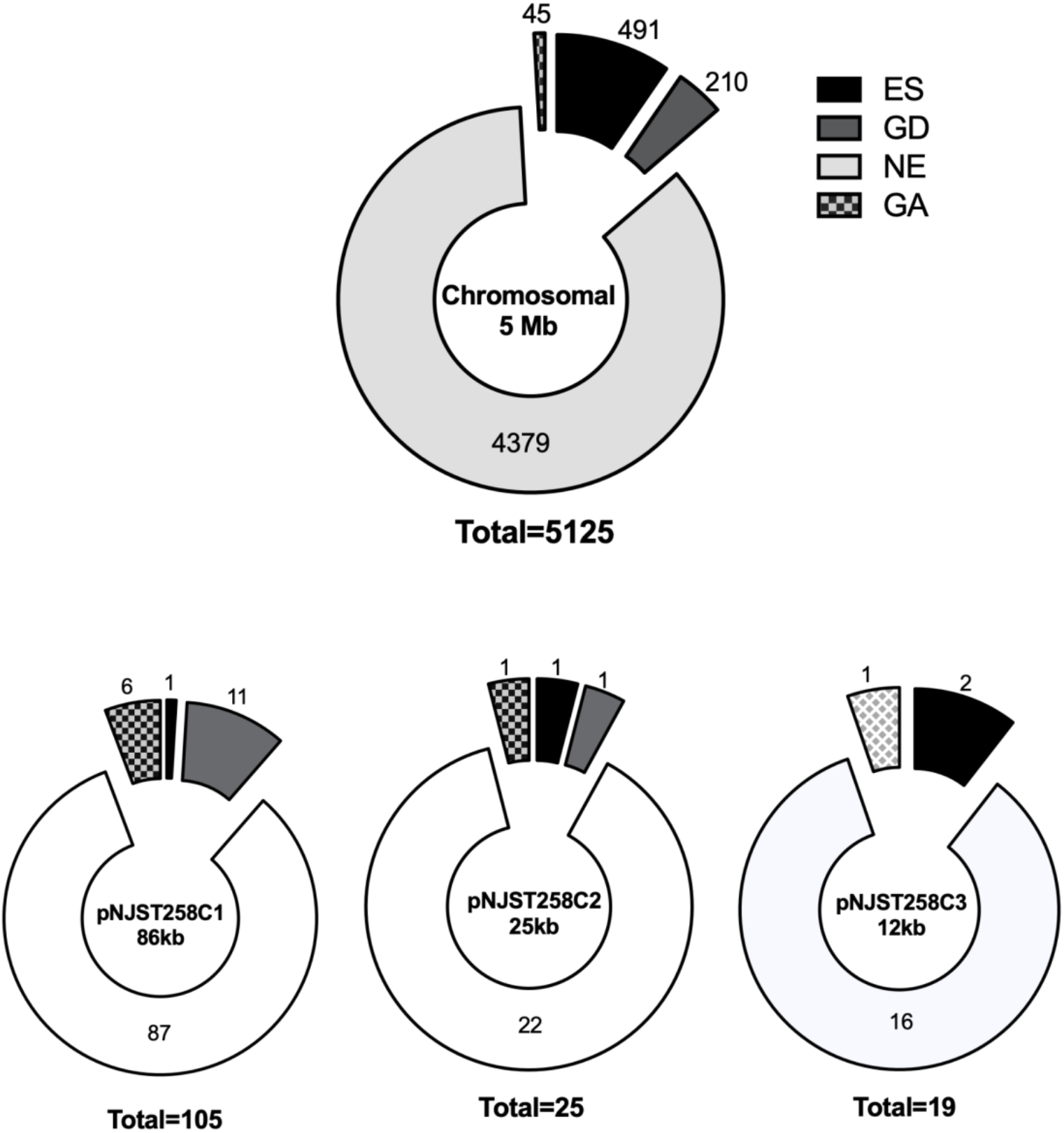
Hidden Markov Model (HMM) analysis and functional categorization of chromosomal and plasmid loci in *Klebsiella pneumoniae* ST258 strain BK30684. **A)** Chromosomal loci are categorized as ES (Essential), NE (Non-Essential), GD (Growth Defect), and GA (Growth Advantage). **(B)** Plasmid loci are categorized as CRT (Critical for plasmid maintenance/stability and/or host fitness), DIS (Dispensable), GD, and GA. Loci excluded due to insufficient coverage are designated N/A. Segment counts are displayed next to each chart.

### Identification of Fitness Determinants Involved in Susceptibility to CFDC

Using the saturated transposon library, we identified genes contributing to the response to CFDC. The ST258 CRKP strain BK30684 library was grown in triplicate in IDMH broth with or without 4μg/mL CFDC for 5 hours. Transposon insertion sites were sequenced, and differential insertion patterns were analyzed using the RESAMPLING module of the TRANSIT software. Genes with significant differences in read counts between IDMH and IDMH+CFDC (*q* ≤ 0.05) were considered conditionally important under drug exposure.

In total, 45 genes were identified (Table 1), including 41 chromosomal genes, 2 on pNJST258C1, and 1 on pNJST258C2. Disruption of genes involved in siderophore uptake (*cirA, tonB*), oxidative stress responses (*cyoA, cyoB, fadE, mdh, ndh*), and nitrogen metabolism (*glnA, glnD*) was associated with increased resistance to CFDC. Disruption of RS27345 and RS27360, located downstream and upstream of the β-lactamase gene *bla*_TEM_ on pNJST258C1, also conferred increased resistance. Notably, multiple genes were involved in peptidoglycan biosynthesis, stability, or regulation, including *mltB, dacA, mrcB, tolA, tolB, ampD, ampG, ldcA, nagZ, lpoB, sltY, mepK*, and *bipP*. Additional genes were linked to cell envelope structure (*wecA, sanA, wzxE, wzyE, rfaL*). Despite the presence of multiple β-lactamases, only *bla*_KPC-3_ encoded on pNJST258C2 showed a significant association with increased CFDC susceptibility.

**Table 1:** Conditionally essential genes in the presence of CFDC. Genes exhibiting significant differences in normalized transposon insertion counts between CFDC-treated and untreated conditions (*q* < 0.05) are listed. Conditional enrichment is expressed as log_2_(CFDC/control), with negative and positive values indicating under- and overrepresentation in CFDC, respectively.

| **Locus** | **Gene** | **Description** | **log<sub>2</sub>FC** |
| --- | --- | --- | --- |
| RS10880 | *tonB* | TonB system transport protein | 4.37 |
| RS21890 | *glnD* | uridylyltransferase/uridylyl-removing protein | 3.37 |
| RS01975 | *fkpA* | FKBP-type peptidyl-prolyl *cis-trans* isomerase | 2.09 |
| RS02750 | *mdh* | malate dehydrogenase | 2.09 |
| RS26305 | *glnA* | glutamate ammonia ligase | 1.86 |
| RS02175 | *purH* | AICAR transformylase / IMP cyclohydrolase | 1.68 |
| RS03355 | *yqjA* | DedA family protein | 1.52 |
| RS20710 | *lon* | endopeptidase La | 1.48 |
| RS08565 | *cirA* | catecholate siderophore receptor CirA | 1.45 |
| RS21615 | *fadE* | acyl-CoA dehydrogenase | 1.35 |
| RS20745 | *cyoA* | cytochrome o ubiquinol oxidase subunit II | 1.31 |
| RS20750 | *cyoB* | cytochrome o ubiquinol oxidase subunit I | 1.3 |
| RS16605 | *ndh* | NAD(P)/FAD-dependent oxidoreductase | 1.29 |
| P1_RS27345 |  | IS6-like element IS26 family transposase | 1.17 |
| P1_RS27360 |  | Tn3-like element Tn3 family transposase | 1.12 |
| RS02670 | *csrD* | RNase E specificity factor CsrD | 1.03 |
| RS25125 | *uvrA* | excinuclease ABC subunit UvrA | -1.04 |
| RS07335 | *ppk1* | polyphosphate kinase 1 | -1.08 |
| RS17535 | *clpA* | ATP-dependent Clp protease ATP-binding subunit | -1.11 |
| RS16620 | *thiK* | thiamine kinase | -1.18 |
| RS25750 | *wecA* | UDP-GlcNAc–Und-P GlcNAc-1-phosphotransferase | -1.32 |
| RS17335 | *mepK* | M15-peptidase | -1.46 |
| RS05650 | *barA* | two-component sensor histidine kinase BarA | -1.47 |
| RS08605 | *sanA* | outer membrane permeability protein SanA | -1.48 |
| RS26005 | *metL* | aspartate kinase/homoserine dehydrogenase | -1.48 |
| RS25710 | *wzxE* | lipid III flippase WzxE | -1.64 |
| RS01215 | *bipP* | M16 metallopeptidase | -1.67 |
| RS06130 | *mltB* | lytic murein transglycosylase B | -1.68 |
| RS03775 | *cpdA* | 3',5'-cyclic-AMP phosphodiesterase | -1.86 |
| RS25700 | *wzyE* | ECA oligosaccharide polymerase | -1.92 |
| RS00740 | *rfaL* | O-antigen ligase family protein | -2.07 |
| RS19070 | *dacA* | D-alanyl-D-alanine carboxypeptidase DacA | -2.32 |
| RS10180 | *dsbB* | disulfide bond formation protein DsbB | -2.39 |
| RS21970 | *mrcB* | bifunctional glycosyl transferase/transpeptidase | -2.43 |
| RS16630 | *ycfL* | YcfL family protein | -2.54 |
| RS18635 | *tolA* | cell envelope integrity protein TolA | -2.64 |
| RS18630 | *tolB* | Tol-Pal system protein TolB | -2.9 |
| RS22280 | *ampD* | 1,6-anhydro-N-acetylmuramyl-L-alanine amidase | -2.9 |
| RS10220 | *ldcA* | muramoyltetrapeptide carboxypeptidase | -2.95 |
| RS16615 | *nagZ* | β-N-acetylhexosaminidase | -2.98 |
| RS20740 | *ampG* | muropeptide MFS transporter AmpG | -3.48 |
| RS16625 | *lpoB* | penicillin-binding protein activator LpoB | -3.68 |
| RS20735 | *yajG* | lipoprotein | -3.77 |
| RS22930 | *sltY* | murein transglycosylase | -4.67 |
| P2_RS26705 | KPC | carbapenem-hydrolyzing class A β-lactamase | -4.68 |

These results align with the known mechanism of CFDC activity, which exploits siderophore-mediated iron uptake, while expanding its genetic context by implicating oxidative stress response, cell envelope functions, and plasmid-associated loci in *K. pneumoniae*.

### Validation of Tn-Seq Results

To validate the genetic determinants identified by Tn-Seq, we constructed targeted deletion mutants representing key functional categories implicated in CFDC response, including peptidoglycan remodeling, outer membrane and enterobacterial common antigen (ECA) biogenesis, iron uptake, nitrogen metabolism, and oxidative stress pathways. We also included strains lacking plasmid-borne β-lactamases to assess their contribution to CFDC susceptibility.

Minimum inhibitory concentrations (MICs) for CFDC, FEP, CAZ were measured for each mutant relative to the parental BK30684 strain (Table 2). Consistent with the Tn-Seq findings, Δ*tonB* and Δ*cirA* showed the strongest CFDC resistance phenotypes, with MIC increases 128-fold and 8-fold, respectively. Δ*glnD* exhibited a moderate (≥4-fold) increase, while the pNJST258C1-cured derivative and the Δ*cyoA* mutant displayed smaller (~2-fold) increases. MICs for FEP and CAZ remained unchanged across these mutants, except for the plasmid-cured strain, which showed a notable decrease in FEP MIC, reflecting loss of *bla*_TEM_ β-lactamase activity.

**Table 2:** Antimicrobial susceptibility of *K. pneumoniae* BK30684 and its deletion derivatives. MICs (μg/mL) for the wild-type BK30684 strain and its deletion mutants were determined by broth microdilution. Dark shading indicates a g2-fold increase in MIC relative to the wild-type (reduced susceptibility), whereas light shading indicates a g2-fold decrease (increased susceptibility). Antibiotics tested: cefiderocol (CFDC), cefepime (FEP), and ceftazidime (CAZ).

| Strain | CFDC | FEP | CAZ |
| --- | --- | --- | --- |
| WT | 0.06 | 32 | 256 |
| $\Delta tonB$ | 8 | 32 | 256 |
| $\Delta glnD$ | 0.25 | 32 | 256 |
| $\Delta cirA$ | 0.5 | 32 | 256 |
| $\Delta cyoA$ | 0.125 | 64 | 256 |
| $\Delta pNJST258C1$ | 0.125 | 4 | 256 |
| $\Delta thiK$ | 0.06 | 32 | 256 |
| $\Delta mepK$ | 0.06 | 32 | 256 |
| $\Delta sanA$ | 0.06 | 32 | 256 |
| $\Delta wzxE$ | 0.03 | 8 | 256 |
| $\Delta bipP$ | 0.06 | 32 | 256 |
| $\Delta mltB$ | 0.06 | 32 | 256 |
| $\Delta wzyE$ | 0.06 | 32 | 256 |
| $\Delta rfaL$ | 0.03 | 32 | 256 |
| $\Delta mrcB$ | 0.03 | 32 | 256 |
| $\Delta ycfL$ | 0.06 | 32 | 256 |
| $\Delta ampD$ | 0.06 | 32 | 256 |
| $\Delta ldcA$ | 0.03 | 32 | 256 |
| $\Delta nagZ$ | 0.06 | 32 | 256 |
| $\Delta ampG$ | 0.03 | 32 | 256 |
| $\Delta lpoB$ | 0.06 | 32 | 256 |
| $\Delta yajG$ | 0.06 | 32 | 256 |
| $\Delta sltY$ | 0.06 | 32 | 256 |
| $\Delta bla_{KPC}$ | 0.03 | >0.25 | >0.25 |
| $\Delta ampG \Delta nagZ$ | 0.06 | 32 | 256 |
| $\Delta mrcB \Delta lpoB$ | 0.03 | 32 | 256 |
| $\Delta ldcA \Delta sltY$ | 0.015 | 16 | 128 |
| $\Delta mltB \Delta sltY$ | 0.06 | 32 | 256 |
| $\Delta mltB \Delta bipP$ | 0.06 | 32 | 256 |
| $\Delta ldcA \Delta bipP$ | 0.03 | 32 | 256 |
| $\Delta rfaL \Delta wzxE$ | 0.015 | 8 | 128 |
| $\Delta rfaL \Delta wzyE$ | 0.03 | 8 | 128 |
| $\Delta wzxE \Delta wzyE$ | 0.015 | 8 | 64 |
| $\Delta wzxE \Delta wzyE \Delta bla_{KPC}$ | 0.0078 | >0.25 | >0.25 |

Given the high redundancy of envelope biogenesis proteins (47,48), we generated single and double knockouts targeting functional modules (49), including ECA biosynthesis, peptidoglycan recycling, transglycosylases, and endopeptidases. Most single-gene deletions did not appreciably alter CFDC MICs; however, a subset (Δ*wzxE*, Δ*ldcA*, Δ*ampG*, Δ*lpoB*, Δ*mrcB, ΔrfaL*) exhibited modest (~2-fold) reductions. Inactivation of multiple genes within the same pathway resulted in moderate reductions in CFDC MICs, consistent with functional overlap. Similarly, the Δ*bla*_KPC_ strain showed a slight increase in CFDC susceptibility, but its MICs for FEP and CAZ dropped dramatically, consistent with the central role of *bla*_KPC_ in resistance to conventional cephalosporins. Importantly, combining deletion of *bla*_KPC_ with disruption of ECA biosynthesis (Δ*wzxE* Δ*wzyE*) resulted in a marked increase in CFDC susceptibility, with an MIC that was 8-fold lower than that of the parental BK30684 strain and lower than either the Δ*bla*_KPC_ or Δ*wzxE* Δ*wzyE* mutants alone. This result indicates that the CFDC tolerance phenotype arises from the combined contributions of multiple determinants.

To further assess these contributions beyond endpoint MIC values, we monitored growth under sub-inhibitory CFDC concentrations (0.25× wild-type MIC) (Figure 2). In addition to mutants displaying MIC changes, several single-gene mutants without detectable MIC shifts (Δ*ampD*, Δ*sltY*, Δ*nagZ*, Δ*mltB*) exhibited prolonged lag phases and reduced growth rates. Notably, double mutants (Δ*wzxE* Δ*wzyE*, Δ*rfaL* Δ*wzyE*, Δ*ldcA* Δ*sltY*) showed pronounced growth defects compared to their respective single-gene knockouts. These findings highlight that simultaneous inactivation of redundant genes significantly exacerbates susceptibility to CFDC. Collectively, these results demonstrate that Tn-Seq uncovers genetic determinants of CFDC tolerance that are functionally critical under stress conditions.

**Figure 2.**
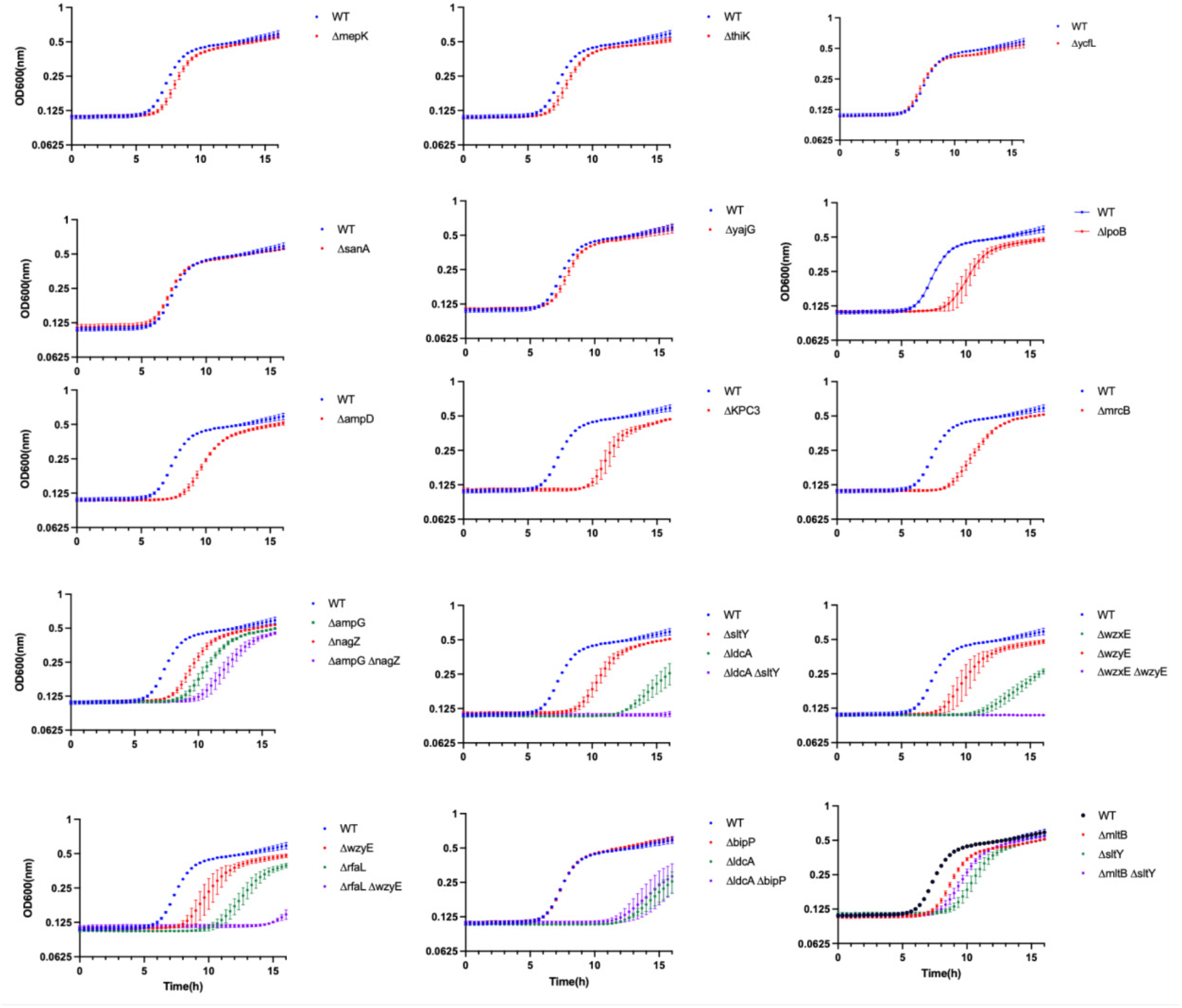
Growth kinetic of *Klebsiella pneumoniae* BK30684 and its deletion derivatives in the presence of CFDC. Wild1type BK30684 and deletion mutants were inoculated in IDMH broth supplemented with CFDC at 0.25× the MIC of the wild type, and growth was monitored by OD600 for 16h. Data represent means ± standard deviation from three independent biological replicates.

### Siderophore transport defects limit the impact of intrinsic resistance determinants

While our Tn-Seq analysis and initial validation confirmed that the abrogation of siderophore uptake (via TonB or CirA disruption) is the primary driver of high-level CFDC resistance, it remains unclear whether this resistance barrier can be lowered by targeting the intrinsic resistome. We hypothesized that compromising cell envelope homeostasis could introduce a secondary vulnerability, rendering Δ*tonB* mutants susceptible to CFDC despite their transport defects. To test this hypothesis, we engineered double mutants by introducing deletions of candidate sensitizing genes, specifically those involved in peptidoglycan metabolism (*ampG, ldcA, mltB-sltY, lpoB*) and ECA biosynthesis (*wzxE*-*wzyE*), into the CFDC-resistant Δ*tonB* background. We also included a Δ*bla*_KPC-3_ mutant to assess the impact of hydrolytic burden in the absence of active transport. These double mutants were evaluated via growth kinetic and MIC assays to determine if intrinsic envelope stress could override the protection conferred by the loss of iron transport (Figure 3).

**Figure 3.**
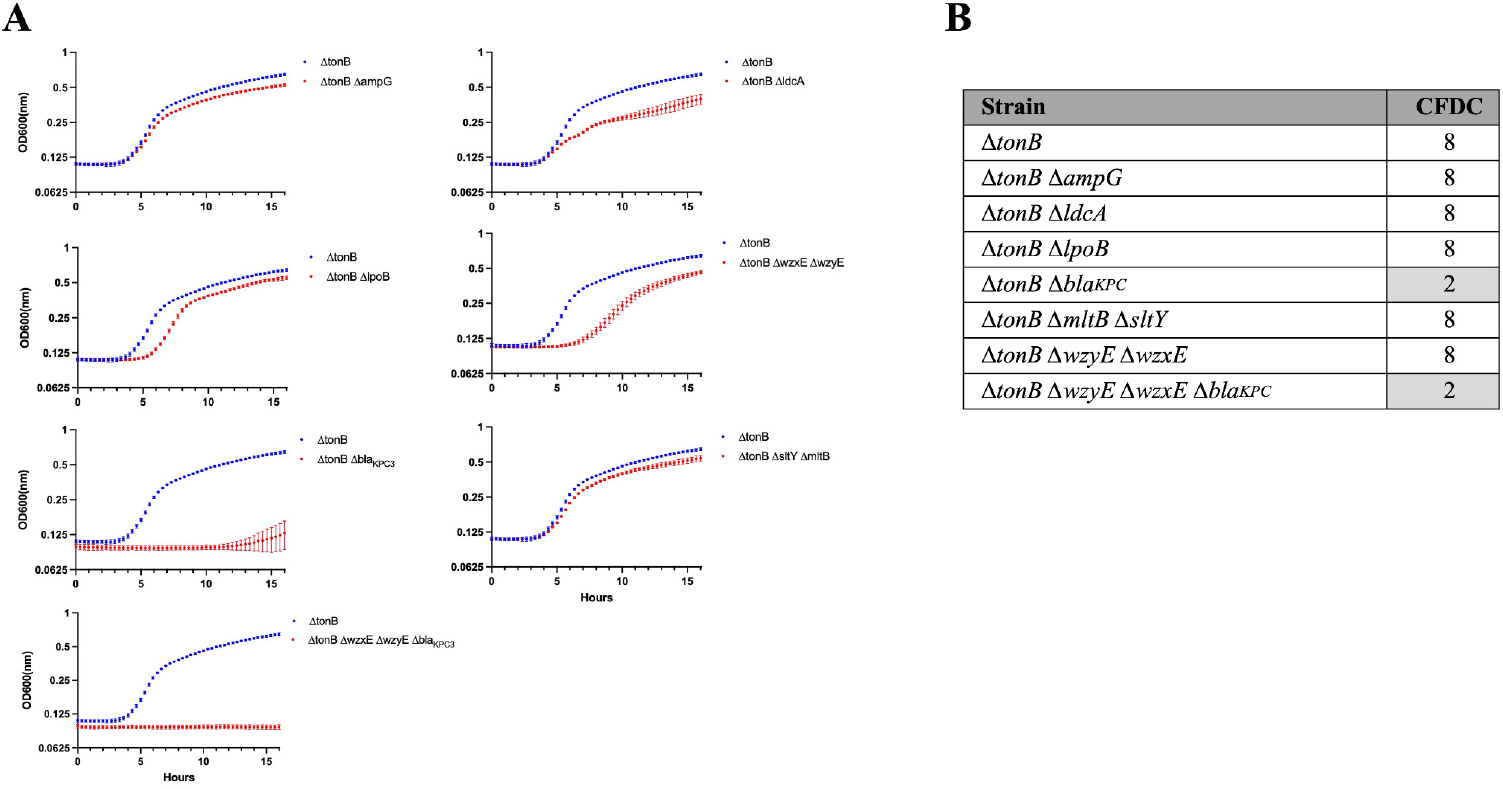
Assessment of susceptibility restoration in a cefiderocol-resistant background. **(A)** Growth kinetic of resensitization target (RT) gene deletions constructed in the Δ*tonB* background were evaluated under 0.25× the MIC of cefiderocol determined for the Δ*tonB* strain. Growth was monitored by OD600 for 16h. Data represent means ± standard deviation from three independent biological replicates. **(B)** CFDC MICs (μg/mL) for &*tonB* strain and its deletion mutants were determined by broth microdilution. Light shading indicates a g2-fold decrease relative &*tonB* parental background.

In contrast to results observed in the wild-type background, deletion of *ampG* or *mltB–sltY* in the Δ*tonB* strain did not significantly alter growth kinetic in the presence of CFDC. Inactivation of *lpoB* or *wzxE– wzyE* resulted in increased lag phases, whereas deletion of *ldcA* produced a modest reduction in overall growth relative to the Δ*tonB* parental strain. However, the magnitude of these growth defects was substantially attenuated compared to that observed in the wild-type background. Notably, deletion of *bla*_KPC-3_ in the Δ*tonB* background resulted in pronounced growth impairment at 0.25× the Δ*tonB* CFDC MIC and produced a 4-fold reduction in MIC compared with the Δ*tonB* parental strain. These findings support a measurable contribution of β-lactamase activity to CFDC resistance, even in the context of compromised siderophore-mediated uptake.

Collectively, these data indicate that while perturbation of cell envelope–associated intrinsic resistance pathways can partially sensitize CFDC-resistant Δ*tonB* mutants, the overall impact of targeting downstream intrinsic resistance determinants is substantially constrained by limited drug uptake.

## Discussion

Cefiderocol (CFDC) was developed in order to circumvent the permeability barriers and β-lactamase mediated resistance mechanisms that limit the efficacy of conventional cephalosporins against multidrug-resistant Gram-negative bacteria (12,14,15). Despite this design, clinical CFDC resistance has been increasingly reported in *Klebsiella pneumoniae*, and surveillance studies have linked reduced susceptibility primarily to defects in siderophore-mediated iron uptake and the emergence of specific β-lactamases, including KPC variants and metallo-β-lactamases such as NDM (13,17–19,21,29,31,33). Nevertheless, these primary determinants do not fully account for the heterogeneous CFDC phenotypes observed among closely related *K. pneumoniae* isolates, nor do they explain differences in bacterial fitness and survival during CFDC exposure. In this study, we employed genome-wide transposon mutagenesis to define the broader genetic landscape underlying CFDC susceptibility in *K. pneumoniae*, revealing a network of interacting loss-of-function determinants that collectively shape bacterial fitness under CFDC exposure.

Our Tn-Seq analysis identified the siderophore-mediated iron uptake machinery as the primary determinant of CFDC susceptibility (18–21,29). Insertions in *tonB* and *cirA* were among the most highly enriched loci under CFDC selection, with targeted deletion mutants exhibiting MIC increases of 128-fold and 8-fold, respectively. The markedly higher resistance observed in the Δ*tonB* mutant relative to Δ*cirA* indicates that, while CirA is the primary portal (18–20), other outer-membrane siderophore receptors likely participate in CFDC uptake. Indeed loss of TonB - the universal energy transducer for outer-membrane siderophore transport-abolishes CirA-mediated CFDC uptake and collapses all TonB-dependent entry routes, resulting in significantly higher resistance (17,26,27,29,50,51).

Beyond uptake, most enriched loci modulated bacterial survival, revealing additional physiological layers that shape CFDC responses. In particular, genes involved in aerobic respiration and redox balance (including *cyoA, cyoB, ndh, mdh*, and *fadE*) were significantly overrepresented during CFDC exposure. Targeted validation of a representative respiratory locus (*cyoA*) confirmed that disruption of aerobic electron transport confers a modest but measurable survival advantage under CFDC challenge. These observations align with emerging evidence that suppression of aerobic respiration and oxidative stress responses can attenuate the bactericidal activity of CFDC and other β-lactams by limiting metabolism-dependent secondary damage (27,35,52,53). Our analysis further implicated nitrogen homeostasis as a determinant of CFDC susceptibility, evidenced by the enrichment of mutants in *glnD* (uridylyltransferase) and *glnA* (glutamine synthetase), two core components of the PII-regulated nitrogen assimilation pathway. Targeted validation of the Δ*glnD* mutant confirmed that disruption of central nitrogen regulation confers a reproducible survival advantage (≥4-fold MIC increase). Notably, this mutant exhibited a marked growth defect, a phenotype indicative of metabolic quiescence. This slow-growth state is well-documented to antagonize the bactericidal activity of cell-wall-active agents (54–57). Alongside *glnD, csrD* was enriched under CFDC selection; recent reports suggest that inactivation of these loci results in capsule-mediated drug exclusion (58). Collectively, these data delineate the loss-of-function determinants underlying CFDC resistance.

Our analysis also identified an intrinsic “sensitization module” comprising loci whose loss increases CFDC susceptibility. The largest component of this module mapped to peptidoglycan biosynthesis, including PBP1-associated synthetic enzymes (*mrcB, lpoB*, and *dacA*) and muropeptide turnover factors (*ampG, ampD, nagZ, mltB*, and *sltY*). The prominence of these loci is consistent with the primary cellular target of CFDC peptidoglycan synthesis via inhibition of PBP3 (22). Proteins involved in peptidoglycan metabolism are highly redundant, with overlapping functions that buffer the impact of individual gene disruptions (47,48). Accordingly, most single deletions resulted in only modest decreases in CFDC MICs. Despite these limited endpoint effects, growth-kinetic assays revealed pronounced fitness defects during CFDC exposure for multiple mutants (e.g., Δ*ldcA, ΔsltY, ΔampG, ΔmrcB, ΔlpoB*, and Δ*ampD*), highlighting their collective role in envelope adaptation during sustained cell-wall stress. This redundancy was further underscored by combinatorial disruptions, such as Δ*ldcA-ΔsltY*, which resulted in substantially greater fitness defects than either single mutation alone. Critically, the sensitizing potential of these intrinsic determinants was hierarchically constrained. Disruption of peptidoglycan-associated pathways in a Δ*tonB* background failed to restore CFDC susceptibility, as the contribution of these genes was markedly attenuated relative to the wild-type background. In contrast, deletion of *bla*_KPC-3_ in the Δ*tonB* background resulted in a significant reduction in CFDC resistance. This pattern is consistent with a model in which reduced CFDC influx in the Δ*tonB* background lowers intracellular drug exposure to levels at which the otherwise limited anti-CFDC activity of KPC β-lactamase becomes sufficient to drive resistance (9,30,59–62). Distinct from the peptidoglycan network, we identified a critical dependency on the ECA biosynthetic pathway, specifically the flippase *wzxE* and the polymerase *wzyE* (63–65). Notably, inactivation of *wzxE* or *wzyE* in a Δ*tonB* background significantly impaired bacterial growth during CFDC exposure, identifying the ECA pathway as a potential vulnerability in high-level CFDC-resistant strains.

Collectively, these findings define a hierarchical framework for CFDC resistance in *K. pneumoniae*. TonB-dependent transport constitutes the primary entry layer. Following CFDC entry, β-lactamase activity provides a secondary layer of resistance, complemented by a distributed network of peptidoglycan remodeling and envelope-stabilizing pathways that support survival during sustained cell-wall stress. Although targeting individual intrinsic resistance mechanisms can partially impair growth in siderophore uptake–deficient backgrounds, such interventions are insufficient to fully restore CFDC susceptibility in the absence of functional uptake. Taken together, this work provides a comprehensive functional genomics framework for CFDC resistance that may inform strategies to preserve the clinical utility of this critical antibiotic.

